# NUDT21 links mitochondrial IPS-1 to RLR-containing stress granules and activates host antiviral defense

**DOI:** 10.1101/2020.04.09.033597

**Authors:** Saeko Aoyama-Ishiwatari, Tomohiko Okazaki, Shun-ichiro Iemura, Tohru Natsume, Yasushi Okada, Yukiko Gotoh

## Abstract

Viral RNA in the cytoplasm of mammalian host cells is recognized by retinoic acid– inducible protein–I (RIG-I)–like receptors (RLRs), which localize to cytoplasmic stress granules (SGs). Activated RLRs associate with the mitochondrial adaptor protein IPS-1, which activates antiviral host defense mechanisms including type I interferon (IFN) induction. It has remained unclear, however, how RLRs in SGs and IPS-1 in the mitochondrial outer membrane associate physically and engage in information transfer. Here we show that NUDT21, an RNA binding protein that regulates alternative transcript polyadenylation, physically associates with IPS-1 and mediates its localization to SGs in response to transfection with poly(I:C), a mimic of viral double-stranded RNA. We found that, despite its well-established function in the nucleus, a fraction of NUDT21 localizes to mitochondria in resting cells and becomes localized to SGs in response to poly(I:C) transfection. NUDT21 was also found to be required for efficient type I IFN induction in response to viral infection. Our results together indicate that NUDT21 links RLRs in SGs to mitochondrial IPS-1 and thereby activates host defense responses to viral infection.

## Introduction

The innate immune system provides the first line of defense against viral infection. The initial step of this defense is detection of “non-self” cues known as pathogen-associated molecular patterns (PAMPs) by specialized sensors, known as pattern recognition receptors (PRRs), in host cells. Recognition of viral PAMPs by PRRs results in the activation of a series of mechanisms to combat viral propagation. In vertebrates, activation of PRRs induces the production of type I interferons (IFNs) such as IFN-α and IFN-β and the subsequent expression of hundreds of IFN-stimulated genes (ISGs) that play a major role in restriction of viral replication within infected cells (1–3).

Retinoic acid–inducible protein–I (RIG-I)–like receptors (RLRs) are a family of PRRs consisting of DEAD box–containing RNA helicases that recognize viral RNA in the cytoplasm. Among RLRs, RIG-I recognizes RNA molecules containing a 5’-triphosphate group as well as relatively short (<100 bp) double-stranded RNAs (dsRNAs), whereas melanoma differentiation–associated gene 5 (MDA5) recognizes relatively long (>2 kb) dsRNAs, with both types of dsRNA being derived from a wide range of RNA viruses (4–6). The binding of RLRs to such viral RNAs triggers their interaction with a key antiviral hub protein, IFN-β promoter stimulator–1 (IPS-1, also known as MAVS, CARDIF, and VISA) (7–10). IPS-1 is anchored to the mitochondrial outer membrane, and activated IPS-1 forms large prionlike aggregates on these organelles (11) that in turn activate transcription factors such as interferon regulatory factor 3 (IRF3), nuclear factor–κB, and activator protein–1, resulting in the expression of type I IFNs (8,12–16). The pivotal role of IPS-1 in antiviral responses is exemplified by the finding that IPS-1–deficient mice are more vulnerable to viral infection than are wild-type (WT) mice (17, 18). The NH2-terminal caspase activation and recruitment domain (CARD) of RLRs is required for the interaction with IPS-1 and was indeed found to be essential for IFN induction (19).

Viral RNAs, RLRs, and RLR-associated proteins have been found to be localized to stress granules (SGs) in virus-infected cells. SGs are membraneless structures that are composed of translation-stalled mRNAs and proteins and in which translation is generally inhibited (20). They are formed in response to various cellular stresses including viral infection. Viral RNA–mediated activation of dsRNA-activated protein kinase (PKR) results in the phosphorylation of eukaryotic initiation factor 2a, translational arrest, and SG nucleation (21). The RNA binding proteins Ras-GAP SH3 domain–binding protein 1 (G3BP1), T cell–restricted intracellular antigen 1 (TIA1), and TIA1-related protein (TIAR) participate in the formation of SGs (22, 23), which is thought to involve liquid-liquid phase separation (24). SGs have been proposed to serve as a platform for RLR recognition of viral RNA and consequent activation of antiviral responses, with such SGs also having been termed antiviral SGs (avSGs) (25). Inhibition of SG formation by depletion of G3BP1 indeed suppressed type I IFN expression in response to infection with influenza A virus or Newcastle disease virus (NDV) (25, 26). The fact that many viral factors interfere with SG formation (27–31) also supports an antiviral function of SGs.

The mitochondrial localization of IPS-1 has been thought to be essential for its function. For instance, deletion of the COOH-terminal transmembrane (TM) domain of IPS-1 abrogated both its mitochondrial localization and the induction of type I IFN (8, 32). Caspase-mediated cleavage of IPS-1 that results in detachment of the TM domain also inactivates IPS-1 function (33, 34). Given the mitochondrial localization of IPS-1, it has remained unclear how RLRs within SGs encounter and activate IPS-1. Of interest in this regard, a fraction of IPS-1 appears to colocalize with TIAR, a marker of SGs, in cells infected with viruses or transfected with polyinosinic-polycytidylic acid [poly(I:C)], a synthetic analog of viral dsRNA (26, 35), suggesting that viral RNA may induce the localization of IPS-1 to SGs and its association with RLRs. The mechanism that might underlie such an effect has remained unknown, however.

We here identify nucleoside diphosphate–linked moiety X (Nudix)–type motif 21 (NUDT21) as an IPS-1 interactor. NUDT21 is an RNA binding protein that constitutes the CFIm complex in nucleus and regulates the choice of polyadenylation site (PAS) by binding to the UGUA element in the 3’ untranslated region (UTR) of transcripts (36–39). NUDT21-mediated alternative polyadenylation has been shown to influence various cell fate decision processes as well as tumorigenesis (40, 41). Virus infection was previously shown to induce genome-wide changes in PAS selection in host transcripts and was also reported to limit a host gene expression through NUDT21(42, 43). Unexpectedly, we found that, whereas NUDT21 is localized mostly to the nucleus, where alternative polyadenylation takes place, a fraction of NUDT21 is associated with mitochondria in resting cells but also localizes to SGs in response to poly(I:C) transfection. Moreover, NUDT21 was found to associate with IPS-1 and to play an important role in its localization to SGs as well as in the efficient induction of IFN expression in response to poly(I:C) transfection or to infection with encephalomyocarditis virus (EMCV). Our results thus suggest an unexpected role for NUDT21 in the recruitment of mitochondrial IPS-1 to SGs and in the consequent promotion of RLR-mediated activation of IPS-1 and IFN induction in virus-infected cells.

## Materials and Methods

### Cell culture and transfection

HeLa S3 and HEK293T cells were maintained in Dulbecco’s modified Eagle’s medium supplemented with 10% fetal bovine serum and 1% penicillin-streptomycin. HEK293T cells were transfected with the use of the GeneJuice Transfection Reagent (Merck Millipore), whereas transfection of HeLa S3 cells with expression vectors was performed with Lipofectamine 2000 (Thermo Fisher Scientific).

### Plasmids and reagents

The plasmid pEF-BOS-FLAG-IPS-1 encoding WT human IPS-1 was kindly provided by M. Yoneyama (Division of Molecular Immunology, Medical Mycology Research Center, Chiba University, Japan). A full-length cDNA for NUDT21 was amplified by PCR from a mouse cDNA library with the primers 5′ -GGCAGATCTATGTCTGTGGTGCCGCCCAA-3′ and 5′ -GGCGAATTCTCAGTTGTATATAAAATTGA-3′ (sense and antisense, respectively), and was subcloned into either the BglII and EcoRI sites of the pCS4 vector or the BamHI and EcoRI sites of pcDNA3.1. Human G3BP1 cDNAs with or without stop codon were amplified by PCR from pN1/G3BP1-iRFP (Okada lab plasmids, Addgene #129339) with the sense primer 5′ -GCCAGATCTATGGTGATGGAGAAGCCTAG-3′ and the antisense primers 5′ -GCCGAATTCGGATCCTTACTGCCGTGGCGC-3′ or 5′ -GCCGAATTCGGATCCCTGCCGTGGCGCAAG-3′, respectively, and were subcloned into the BamHI and EcoRI sites of pcDNA3. Myc epitope–tagged full-length human IPS-1 and Myc–IPS-1(ΔTM) cDNAs were amplified by PCR from a Myc–IPS-1 expression vector described previously (44) with the sense primer 5′ -ACTGCGGCCGCATGAGCAAAAGCTCATTT-3′ and the antisense primers 5′ -GCCTCTAGACTAGTGCAGACGCCGCCG-3′ or 5′ -GCCTCTAGACTAGTCTACCTGGGATGCCA-3, respectively, and were subcloned into the NotI and XbaI sites of either pcDNA3 or the G3BP1 expression vector. Myc– IPS-1 (amino acids 1–98), Myc–IPS-1 (amino acids 99–200) and Myc– IPS-1 (amino acids 201–540) cDNAs were amplified by PCR from a Myc–IPS-1 expression vector with the sense primer 5′ -GGCAGATCTATGCCGTTTGCTGAAGACAAGACC-3′ and the antisense primer 5′ -GGCAGATCTCTACCGAGGCTGGTAGCTCTGGT-3′, the sense primer 5′ -GGCAGATCTACCTCGGACCGTCCCCCAGAC-3′ and the antisense primer 5′ -GGCAGATCTCTATGTGTCCTGCTCCTGATGCCCGCT-3′, or the sense primer 5′ -GGCAGATCTATGGAACTGGGCAGTACCCACACAGCA-3′ and the antisense primer 5′ -GGCAGATCTCTAGTGCAGACGCCGCCGGT-3′, respectively, and were subcloned into the BglII and EcoRI sites of the pCS4 vector. Poly(I:C) was obtained from GE Healthcare and was introduced into cells by transfection with Lipofectamine 2000 (Thermo Fisher Scientific).

### Antibodies

Antibodies to NUDT21 were obtained from Proteintech; those to Myc (9E10), to p38, to TIAR, and to TOMM20 were from Santa Cruz Biotechnology; those to IPS-1 for immunostaining as well as those to phospho-p38, to phospho-JNK, to cleaved caspase-3, to cleaved PARP, and to phospho-IRF3 were from Cell Signaling Technology; those to FLAG (M2) were from Sigma; those to HA were from Roche; those to IPS-1 for immunoblot analysis were from Abcam; those to cytochrome c were from BD Pharmingen; and those to CFIm68 were from Bethyl Laboratories. Antibodies to RIG-I were kindly provided by M. Yoneyama (Chiba University).

### RNA interference

Knockdown of NUDT21 was achieved with the use of Stealth RNA interference (Thermo Fisher Scientific). HeLa S3 cells were transfected with siRNA oligonucleotides with the use of the Lipofectamine RNAiMAX reagent (Thermo Fisher Scientific) and were used for experiments after incubation for 72 h. The siRNA sequences were 5′ -UGAACCUCCU-CAGUAUCCAU-AUAUU-3′ and 5′ -AAUAUAUGGA-UACUGAGGAG-GUUCA-3′ for NUDT21, and a negative control siRNA (Thermo Fisher Scientific, catalog no. 12935300) was also used.

### Mass spectrometry

Liquid chromatography and tandem mass spectrometry were performed as previously described (45).

### Immunoblot analysis

Immunoblot analysis was performed as described previously (44). In brief, cells were lysed with a solution containing 20 mM Tris-HCl (pH 7.5), 150 mM NaCl, 10 mM β-glycerophosphate, 5 mM EGTA, 1 mM Na4P2O7, 5 mM NaF, 0.5% Triton X-100, 1 mM Na3VO4, 1 mM dithiothreitol, and aprotinin (1 mg/ml), cell lysates were fractionated by SDS-polyacrylamide gel electrophoresis on a 10% gel, and the separated proteins were transferred to a polyvinylidene difluoride membrane. The membrane was incubated first with primary antibodies for 24 h at 4°C and then with horseradish peroxidase– conjugated secondary antibodies (GE Healthcare) for 1 h at room temperature. After a wash with a solution containing 50 mM Tris-HCl (pH 8.0), 150 mM NaCl, and 0.05% Tween 20, the membrane was processed for detection of peroxidase activity with chemiluminescence reagents and an Image Quant LAS4000 instrument (GE Healthcare).

### Co-immunoprecipitation analysis

HEK293T cells harvested 20 h after transfection with plasmids encoding FLAG–IPS-1 and Myc-NUDT21 were lysed with a solution containing 20 mM Tris-HCl (pH 7.5), 150 mM NaCl, 10 mM β-glycerophosphate, 5 mM EGTA, 1 mM Na4P2O7, 5 mM NaF, 0.2% Triton X-100, 1 mM Na3VO4, 1 mM dithiothreitol, and aprotinin (1 mg/ml). The cell lysates were incubated with rotation at 4°C first for 45 min with antibodies to FLAG and then for 45 min with protein A–Sepharose (GE Healthcare). The resulting immunoprecipitates were subjected to immunoblot analysis.

### Immunofluorescence microscopy

Cells were fixed with 4% formaldehyde for 10 min at 37°C, permeabilized with 0.2% Triton X-100 in phosphate-buffered saline (PBS) for 5 min, and incubated for 24 h in PBS containing 2% fetal bovine serum and 2 % bovine serum albumin (blocking buffer). They were then exposed at room temperature first for 1 h to primary antibodies in blocking buffer and then for 1 h to Alexa Fluor–conjugated secondary antibodies (Thermo Fisher Scientific) and Hoechst 33342 in blocking buffer. Moviol were used as mounting medium. Images were acquired with a TCS SP5 confocal microscope (Leica) and were processed with Image J (NIH).

### Colocalization analysis and quantification of SG volume

Manders M1 and M2 coefficients for colocalization were calculated with Coloc 2 of Fiji. A statistical significance test was derived by Costes (46). For the experiment shown in Figure 2C, samples were prepared in the same way as for immunofluorescence analysis described above, with the exceptions that ProLong Diamond (Thermo Fisher Scientific) was used as mounting medium and that images were acquired with a TCS SP8 confocal microscope (Leica). Three-dimensional images were acquired in order to meet the Nyquist condition (pixel size of 40.6 nm for *x* and *y*, and of 100 nm for *z*) and were deconvoluted with Huygens software (Scientific Volume Imaging). Manders M1 and M2 for the colocalization of IPS-1 and TOMM20 were then calculated. For the experiment shown in Figure 2E, samples were again prepared in the same way as for immunofluorescence analysis with the exception that ProLong Diamond (Thermo Fisher Scientific) was used as mounting medium. Three-dimensional images were acquired in order to match the Nyquist condition (pixel size of 50 nm for *x* and *y*, and of 100 nm for *z*) and were deconvoluted with Huygens software (Scientific Volume Imaging). Manders M1 coefficient for the colocalization of IPS-1 and TIAR was then calculated. The SG volume per cell was measured with the 3D object counter (threshold, 25; size filter, 1) of Huygens software in images for which the background intensity of the cytosol had been subtracted.

**Figure 1.**
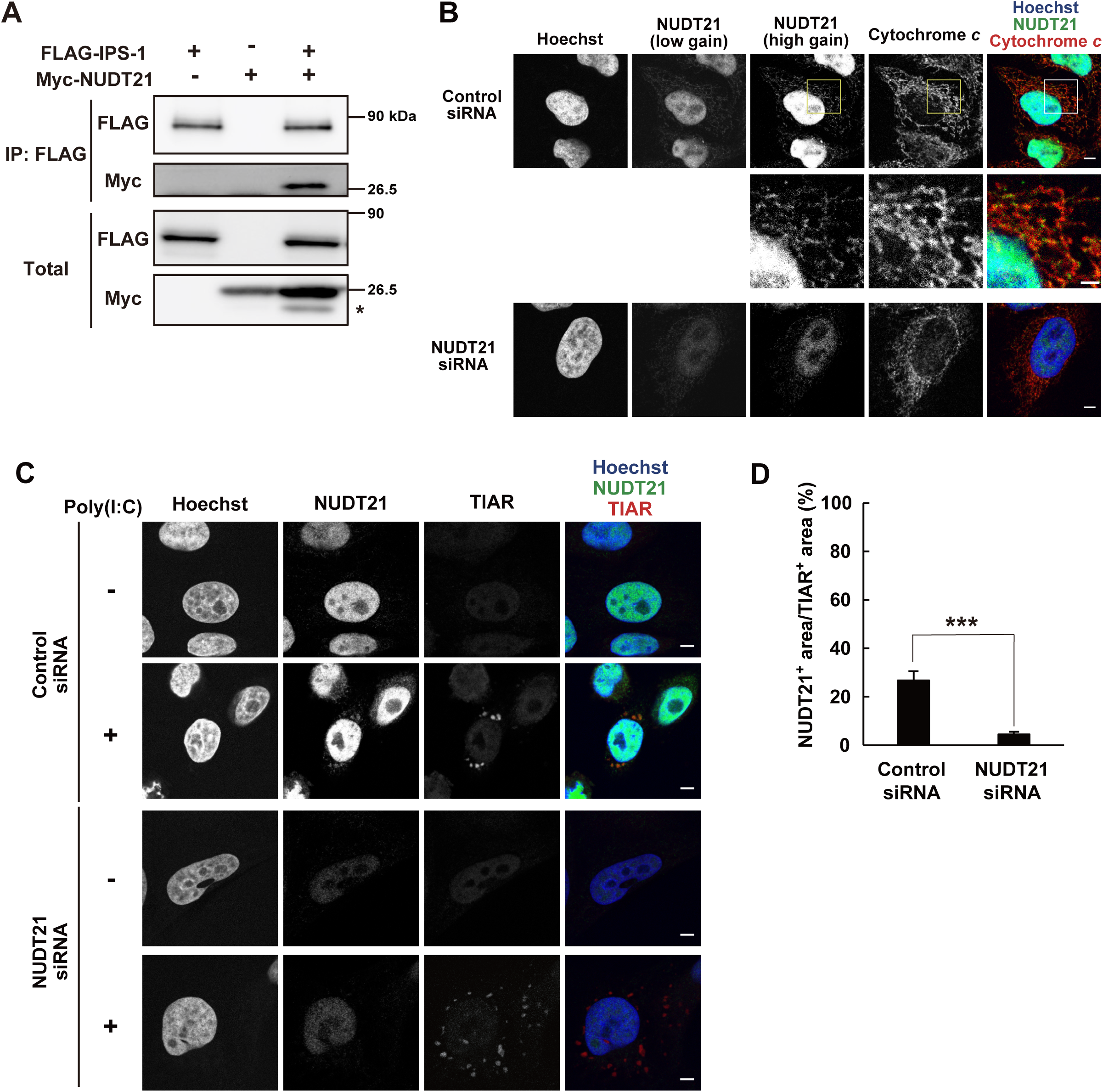
A fraction of NUDT21 localizes to mitochondria in resting cells and contributes to SGs in the cytoplasm in response to poly(I:C) transfection. (A) Extracts of HEK293T cells transiently transfected with expression vectors for FLAG-tagged IPS1 and Myc-tagged NUDT21 were subjected to immunoprecipitation (IP) with antibodies to FLAG, and the resulting precipitates as well as the original cell extracts (Total) were subjected to immunoblot analysis with antibodies to FLAG and to Myc. The asterisk indicates a nonspecific band. Data are representative of three independent experiments. (B) Immunofluorescence analysis of HeLa S3 cells transfected with control or NUDT21 siRNAs. The cells were stained with antibodies to NUDT21 and to cytochrome c as well as with Hoechst 33342. The boxed regions in the upper panels are shown at higher magnification in the corresponding middle panels. Scale bars, 5 μm or 2 μm (higher magnification images). Data are representative of three independent experiments. (C) Immunofluorescence analysis of control and NUDT21-depleted HeLa S3 cells at 6 h after transfection with poly(I:C) (0.25 μg/ml) or mock transfection. The cells were stained with antibodies to NUDT21 and to TIAR as well as with Hoechst 33342. Scale bars, 5 μm. Data are representative of three independent experiments. (D) Quantification of the NUDT21-positive area of TIAR-positive foci per cell in images similar to those in (C). Data are means ± s.e.m. from three independent experiments (control, n = 16; NUDT21 knockdown, n = 13). \*\*\**P* < 0.005 (Student’s *t* test).

**Figure 2.**
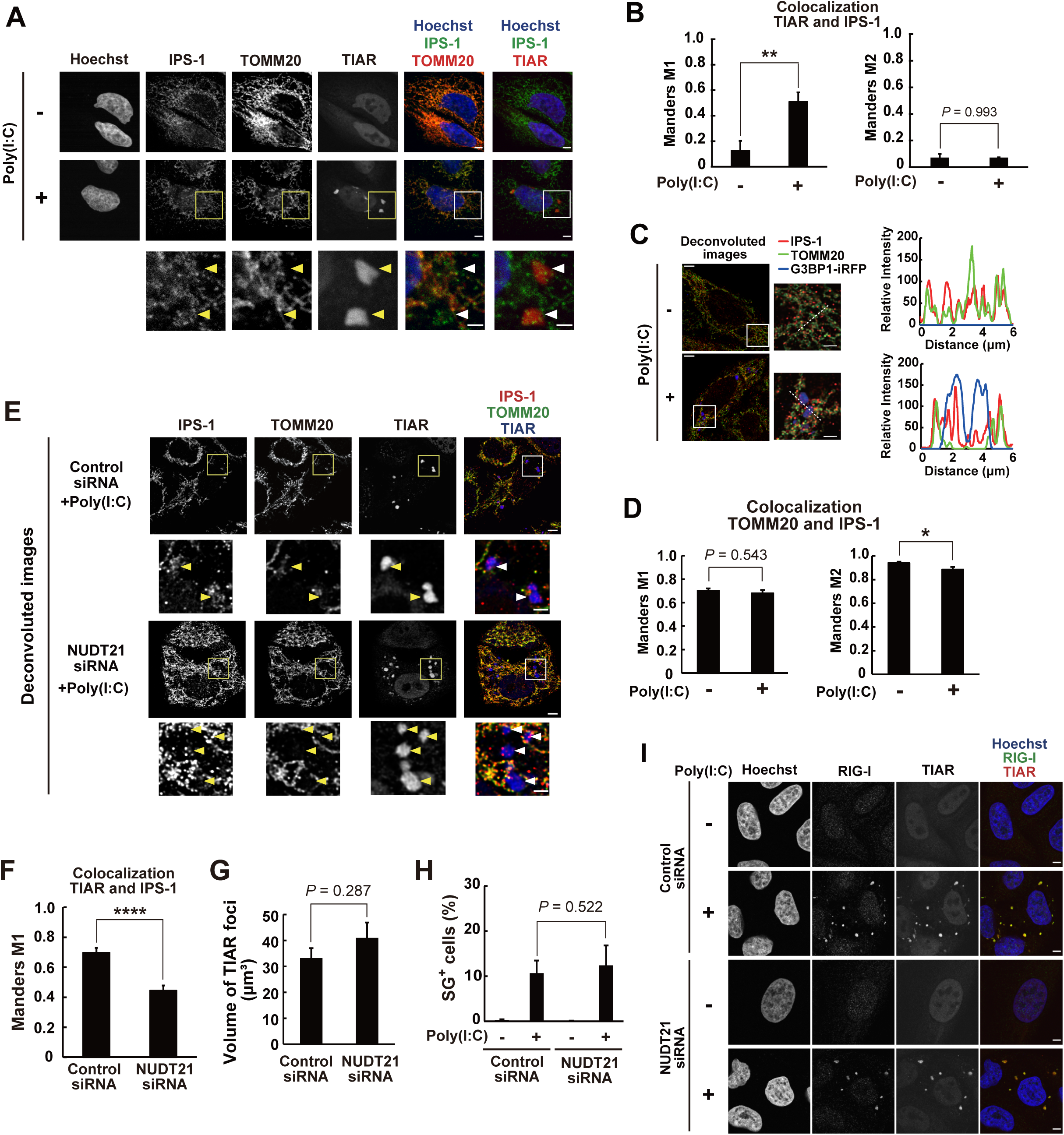
NUDT21 knockdown impairs the localization of IPS-1 to SGs. (A) Immunofluorescence analysis of HeLa S3 cells at 6 h after transfection with poly(I:C) (0.25 μg/ml) or mock transfection. The cells were stained with antibodies to IPS-1, to TOMM20, and to TIAR as well as with Hoechst 33342. The boxed regions of the middle panels are shown at higher magnification in the corresponding lower panels. Arrowheads indicate SGs. Scale bars, 5 μm or 2 μm (higher magnification images). Data are representative of three independent experiments. (B) Colocalization (Manders M1 and M2) between IPS-1 and TIAR was determined for images as in (A). Manders M1 is the sum of TIAR signal intensity overlapping with IPS-1 versus total TIAR signal intensity, whereas Manders M2 is the sum of IPS-1 signal intensity overlapping with TIAR versus total IPS-1 signal intensity. Data are means ± s.e.m. from three independent experiments (mock, n = 5; poly(I:C), n = 11) \*\**P* < 0.01 (Student’s *t* test). (C) HeLa S3 cells transiently expressing iRFP-fused G3BP1 were subjected to immunofluorescence staining with antibodies to IPS-1 and to TOMM20 at 6 h after transfection with poly(I:C) (0.25 μg/ml) or mock transfection. The fluorescence of iRFP was monitored directly. The *z*-stack images were deconvoluted with Huygens software. The boxed regions of the left images are shown at higher magnification in those on the right. Scale bars, 5 μm or 2 μm (higher magnification images). Line-scan analysis of the relative fluorescence intensity of IPS-1, TOMM20, and G3BP1-iRFP along the white broken lines (6 μm) in the enlarged images is also shown. Data are representative of three independent experiments. (D) Colocalization (Manders M1 and M2) between IPS-1 and TOMM20 was determined for images as in (C). Manders M1 is the sum of IPS-1 signal intensity overlapping with TOMM20 versus total IPS-1 signal intensity, and Manders M2 is the sum of TOMM20 signal intensity overlapping with IPS-1 versus total TOMM20 signal intensity. Data are means ± s.e.m. from three independent experiments (mock, n = 12; poly(I:C), n = 18). \**P* < 0.05 (Student’s *t* test). (E) Immunofluorescence analysis of HeLa S3 cells expressing control or NUDT21 siRNAs at 6 h after transfection with poly(I:C) (0.25 μg/ml). The cells were stained with antibodies to IPS-1, to TOMM20, and to TIAR. The *z*-stack images were deconvoluted with Huygens software. The boxed region of the upper image of each pair is shown at higher magnification in the lower image. Arrowheads indicate SGs. Scale bars, 5 μm or 2 μm (higher magnification images). Data are representative of three independent experiments. (F) Colocalization (Manders M1, sum of TIAR signal intensity overlapping with IPS-1 versus total TIAR signal intensity) between IPS-1 and TIAR was determined for images as in (E). Data are means ± s.e.m. of three independent experiments (control, n = 10; NUDT21 knockdown, n = 10). \*\*\*\**P* < 0.001 (Student’s *t* test). (G) Quantification of SG volume per cell for images as in (E). Data are means ± s.e.m. of three independent experiments (control, n = 10; NUDT21 knockdown, n = 10). The *P* value was determined with Student’s *t* test. (H) Quantification of SG-containing cells. HeLa S3 cells expressing control or NUDT21 siRNAs were subjected to immunofluorescence staining for TIAR in order to determine the proportion of SG-containing cells at 6 h after transfection with poly(I:C) (0.25 μg/ml) or mock transfection. Data are means ± s.e.m. of three independent experiments. The *P* value was determined with Student’s *t* test. (I) Immunofluorescence analysis of control and NUDT21-knockdown HeLa S3 cells at 6 h after transfection with poly(I:C) (0.25 μg/ml) or mock transfection. The cells were stained with antibodies to RIG-I and to TIAR as well as with Hoechst 33342. Scale bars, 5 μm. Data are representative of three independent experiments.

### RT and real-time PCR analysis

Total RNA was obtained from cells with the use of RNAiso (TaKaRa). RT was performed with 1 μg of total RNA and ReverTra Ace qPCR RT Master Mix with gDNA Remover (Toyobo). The resulting cDNA was subjected to real-time PCR analysis in a Roche LightCycler with the use of a KAPA SYBR Fast qPCR Kit (Nippon Genetics). The abundance of each target mRNA was normalized by that glyceraldehyde-3-phosphate dehydrogenase (GAPDH) mRNA. The PCR primers (sense and antisense, respectively) were 5′ -ACTCCTCCACCTTTGACGCT-3′ and 5′ -TCCTCTTGTGCTCTTGCTGG-3′ for human GAPDH, 5′ -CTGGCTGGAATGAGACTATTGTT-3′ and 5′ -CTTCAGTTTCGGAGGTAACCTG-3′ for human IFN-β, 5′ -AACCTGAACCTTCCAAAGATGG-3′ and 5′ -TCTGGCTTGTTCCTCACTACT-3′ for human IL-6, and 5′ -ATGAGCACTGAAAGCATGATCC-3′ and 5′ -GAGGGCTGATTAGAGAGAGGTC-3′ for human TNF-α.

### Reporter gene analysis

HEK293T cells seeded in 24-well plates (7.5 × 10^4^ cells/well) were transiently transfected for 20 h with 8 ng of a reporter plasmid encoding *Renilla* luciferase under the control of the human IFN-β gene promoter (p-125-RLuc) together with 5 ng of an expression plasmid encoding FLAG-tagged full-length IPS-1 and either 50 or 500 ng of a plasmid for Myc-tagged IPS-1(ΔTM). The activity of *Renilla* luciferase in total cell lysates was measured with the use of a Dual-Luciferase Reporter Assay System (Promega) and was normalized by that of firefly luciferase derived from 80 ng of a control plasmid.

### Virus infection

EMCV were kindly provided by M. Yoneyama (Chiba University). The cells were incubated in culture medium with 0.1 (+) or 0.2 (++) PFU of EMCV for 2 h, and then replaced in EMCV-free culture medium for 10 h.

### Statistical analysis

Quantitative data are presented as means ± s.e.m. or ± s.d. and were compared with Student’s *t* test. A *P* value of <0.05 was considered statistically significant.

## Results

### NUDT21 forms a complex with IPS-1

In a search for regulators of IPS-1, we performed co-immunoprecipitation analysis to identify IPS-1–associated molecules. FLAG epitope–tagged IPS-1 expressed in HEK293T cells was thus immunoprecipitated with antibodies to FLAG, and the resultant precipitates were analyzed by a highly sensitive direct nanoflow liquid chromatography–tandem mass spectrometry system. We identified NUDT21 among the proteins that coprecipitated with FLAG–IPS-1. To confirm this result, we performed co-immunoprecipitation analysis with HEK293T cells expressing FLAG–IPS-1 and Myc epitope–tagged NUDT21. We found that Myc-NUDT21 coprecipitated with FLAG– IPS-1 (Figure 1A), suggesting that NUDT21 indeed forms a complex with IPS-1.

To examine which domains of IPS-1 interact with NUDT21, we generated Myc-tagged fragments of human IPS-1 that contain either CARD (amino acids 1–98), the proline-rich domain (amino acids 99–200), or the COOH-terminal domain including the TM domain (amino acids 201–540). We found that hemagglutinin epitope (HA)– tagged NUDT21 coprecipitated with Myc–IPS-1 fragments containing CARD or the proline-rich domain, but not with that containing the COOH-terminal domain (Figure S1). These results suggested that IPS-1 associates with NUDT21 through its NH2-terminal domains including CARD and the proline-rich domain.

### A fraction of NUDT21 associates with mitochondria in resting cells and localizes to SGs on exposure to cytoplasmic dsRNA

These results suggestive of a physical interaction between IPS-1 and NUDT21 were unexpected, given the previously described localization of NUDT21 in the nucleus and that of IPS-1 to the mitochondrial outer membrane (8). Immunofluorescence analysis of HeLa S3 cells indeed revealed that most NUDT21 was localized to the nucleus (Figure 1B). However, we detected a fraction of the NUDT21 signals in the cytoplasm (Figure 1B). Knockdown of NUDT21 by transfection of the cells with a small interfering RNA (siRNA) resulted in a reduction in these cytoplasmic and nuclear signals, supporting their specificity for NUDT21 (Figure 1B). The NUDT21 signals in the cytoplasm overlapped with those for the mitochondrial marker cytochrome c (Figure 1B). By contrast, CFIm68, a cofactor of NUDT21 in its nuclear function, did not appear to localize to mitochondria (Figure S2). Together, these results suggested that a fraction of NUDT21 localizes to mitochondria, where it may associate with IPS-1.

We next transfected HeLa S3 cells with the viral dsRNA mimic poly(I:C) to examine its possible effect on the intracellular distribution of NUDT21. NUDT21 immunoreactivity was detected as granulelike aggregates in the cytoplasm at 6 h after poly(I:C) transfection, and these granulelike signals were again attenuated by NUDT21 knockdown (Figure 1C). Moreover, we found that these cytoplasmic signals of NUDT21 colocalized with the SG marker TIAR (Figure 1C). Indeed, 26.89 ± 3.65% of the area of TIAR-positive foci overlapped with NUDT21 immunoreactivity (Figure 1D), suggesting that NUDT21 becomes localized to SGs in the presence of cytoplasmic dsRNA and consequent activation of the RLR pathway.

### NUDT21 plays an important role in IPS-1 localization to SGs

The interaction of NUDT21 with IPS-1 as well as its mitochondrial localization and its appearance at SGs in response to dsRNA exposure prompted us to examine whether NUDT21 regulates the localization of IPS-1. Consistent with previous studies (35, 26), we found that a fraction of IPS-1 colocalized with TIAR in cells transfected with poly(I:C), whereas most IPS-1 appeared to colocalize with the mitochondrial import receptor subunit TOM20 homolog (TOMM20) in both control and poly(I:C)-transfected HeLa S3 cells (Figure 2A). We calculated Manders coefficients for the colocalization of IPS-1 and TIAR (47). Manders M1 (sum of TIAR signal intensity overlapping with IPS-1 versus total TIAR signal intensity) was significantly increased by poly(I:C) transfection (Figure 2B), whereas Manders M2 (sum of IPS-1 signal intensity overlapping with TIAR versus total IPS-1 signal intensity) did not differ significantly between cells with or without poly(I:C) exposure (Figure 2B). These results suggested that a fraction of IPS-1 becomes localized to SGs in response to the presence of cytoplasmic dsRNA. We also found the 34.21 ± 5.88% of the area of TIAR-positive foci overlapped with IPS-1 at 6 h after poly(I:C) transfection, indicating that IPS-1 is not distributed to all SGs. To examine IPS-1 localization at a higher resolution, we stained endogenous IPS-1 and TOMM20 at 6 h after poly(I:C) transfection in HeLa S3 cells expressing a fusion protein of G3BP1 and near-infrared fluorescent protein (iRFP) and obtained *z*-stack images deconvoluted with Huygens software. A fraction of IPS-1 signals was found to localize to G3BP1-iRFP foci in poly(I:C)-transfected cells, whereas TOMM20 was detected exclusively outside of G3BP1-iRFP foci (Figure 2C).

We then examined whether poly(I:C) transfection affects the colocalization of IPS-1 and TOMM20. Although Manders M1 (sum of IPS-1 signal intensity overlapping with TOMM20 versus total IPS-1 signal intensity) was not significantly altered by poly(I:C) transfection, Manders M2 (sum of TOMM20 signal intensity overlapping with IPS-1 versus total TOMM20 signal intensity) was slightly but significantly reduced in cells exposed to poly(I:C) (Figure 2D), suggesting that a fraction of IPS-1 separates from TOMM20 in response to poly(I:C) stimulation. Together, these results indicated that a fraction of IPS-1, but not of another mitochondrial protein (TOMM20), becomes localized to SGs in response to the presence of cytoplasmic dsRNA.

We next examined the effect of NUDT21 on the distribution of IPS-1 in cells transfected with poly(I:C). We thus calculated Manders coefficients for IPS-1 and TIAR in *z*-stack images deconvoluted with Huygens software and found that Manders M1 (sum of TIAR signal intensity overlapping with IPS-1 versus total TIAR signal intensity) was markedly reduced by NUDT21 knockdown (Figure 2E, F). Importantly, NUDT21 knockdown did not affect the overall (mitochondrial) distribution of IPS-1 in control (nontransfected) cells (Figure S3), indicating that the effect of NUDT21 on the localization of IPS-1 was stimulus dependent. By contrast, NUDT21 knockdown did not significantly alter the volume of TIAR-positive foci (SGs) per cell (Figure 2G) or the abundance of IPS-1 (Figure S4). We also found that NUDT21 knockdown did not significantly affect the ratio of SG-containing cells (Figure 2H) or RIG-I accumulation at SGs in response to poly(I:C) transfection (Figure 2I). Collectively, these results suggested that NUDT21 plays a role in IPS-1 localization to SGs in response to the presence of cytoplasmic dsRNA, whereas it does not overtly affect SG formation or RIG-I accumulation at SGs.

### NUDT21 is required for antiviral responses induced by cytoplasmic dsRNA

The interaction between RLRs and IPS-1 triggers activation of downstream signaling pathways in response to virus infection (7–10). Given that our data implicated NUDT21 in localization of IPS-1 to SGs, which contain RLRs, we next asked whether NUDT21 contributes to antiviral responses mediated by RLRs and IPS-1 such as the induction of IFN-β and the proinflammatory cytokines interleukin-6 (IL-6) and tumor necrosis factor–α (TNF-α) in HeLa S3 cells transfected with poly(I:C). We found that NUDT21 knockdown markedly suppressed the increase in the amount of IFN-β mRNA induced by poly(I:C) stimulation (Figure 3A). The up-regulation of IL-6 and TNF-α mRNAs in response to poly(I:C) transfection was also significantly attenuated by depletion of NUDT21 (Figure 3B, C). These results thus suggested that NUDT21 is necessary for the efficient induction of IFN-β and proinflammatory cytokines in response to the presence of cytoplasmic dsRNA.

**Figure 3.**
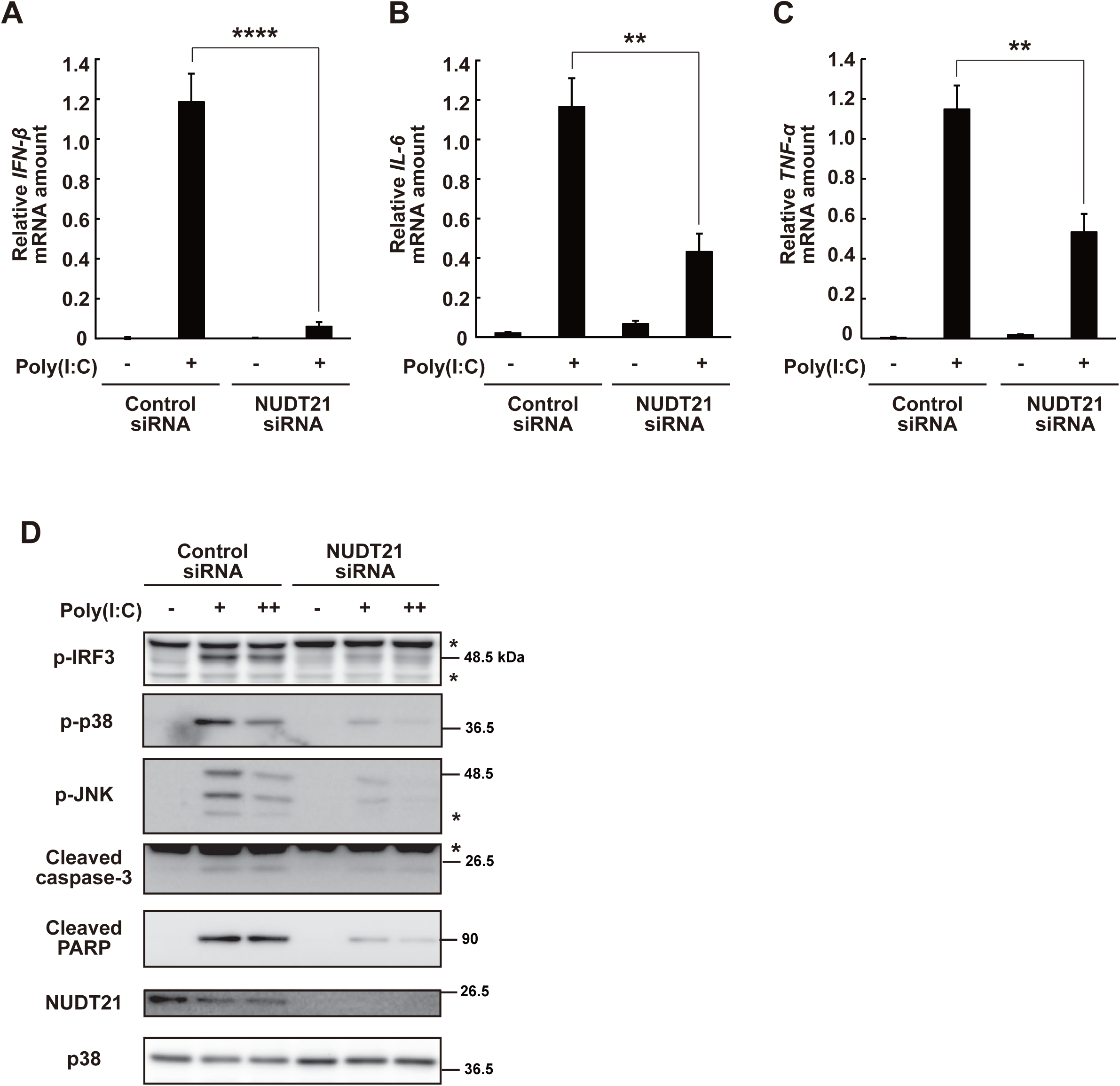
NUDT21 plays a key role in the induction of type I IFN and caspase activation in response to poly(I:C) transfection. (A–C) Quantitative reverse transcription and polymerase chain reaction (RT-PCR) analysis of IFN-β, IL-6, and TNF-α mRNAs, respectively, in HeLa S3 cells transiently expressing control or NUDT21 siRNAs at 9 h after transfection with poly(I:C) (0.25 μg/ml) or mock transfection. Data are expressed relative to the corresponding amount of GAPDH mRNA and are means ± s.e.m. from four independent experiments. \*\**P* < 0.01, \*\*\*\**P* < 0.001 (Student’s *t* test). (D) Immunoblot analysis of phosphorylated (p-) forms of IRF3, p38 MAPK, and JNK as well as of cleaved forms of caspase-3 and PARP in control or NUDT21-knockdwon HeLa S3 cells at 9 h after transfection with poly(I:C) [0.25 (+) or 2.5 (++) μg/ml] or mock transfection. The amounts of NUDT21 and total p38 were examined as controls for knockdown efficiency and protein loading, respectively. Asterisks indicate nonspecific bands. Data are representative of three independent experiments.

We next examined whether NUDT21 mediates activation of the transcription factor IRF3 or the mitogen-activated protein kinases (MAPKs) p38 and c-Jun NH2-terminal kinase (JNK), all of which are essential factors for activation of the promoters of IFN-β and inflammatory cytokine genes in response to cytoplasmic dsRNA (15, 48). Knockdown of NUDT21 suppressed the increase in the abundance of phosphorylated (activated) forms of IRF3 as well as of p38 and JNK induced by poly(I:C) transfection (Figure 3D), suggesting that NUDT21 promotes the activation of these signaling molecules that is essential for the transcription of IFN-β and inflammatory cytokine genes. In addition, NUDT21 knockdown suppressed the cleavage of caspase-3 and poly(ADP-ribose) polymerase (PARP) induced by poly(I:C) transfection (Figure 3D), suggesting that NUDT21 also plays a key role in the induction of caspase activation, and perhaps cell death, in response to the presence of cytoplasmic dsRNA. Together, these results indicated that NUDT21 mediates antiviral cellular responses induced by the presence of cytoplasmic dsRNA.

### NUDT21 mediates type I IFN induction in response to virus infection

To investigate further the role of NUDT21 in antiviral responses triggered by RLRs and IPS-1, we asked whether NUDT21 knockdown affects type I IFN induction in cells infected with viruses. To this end, we studied EMCV, whose RNAs released into the cytoplasm are recognized predominantly by the RLRs MDA5 (6). We found that NUDT21 knockdown significantly suppressed the EMCV-induced phosphorylation of IRF3 and increase in IFN-β mRNA abundance (Figure 4A, B). Together, these results implicated NUDT21 in the optimal induction of IFN-β in response to viral infection.

**Figure 4.**
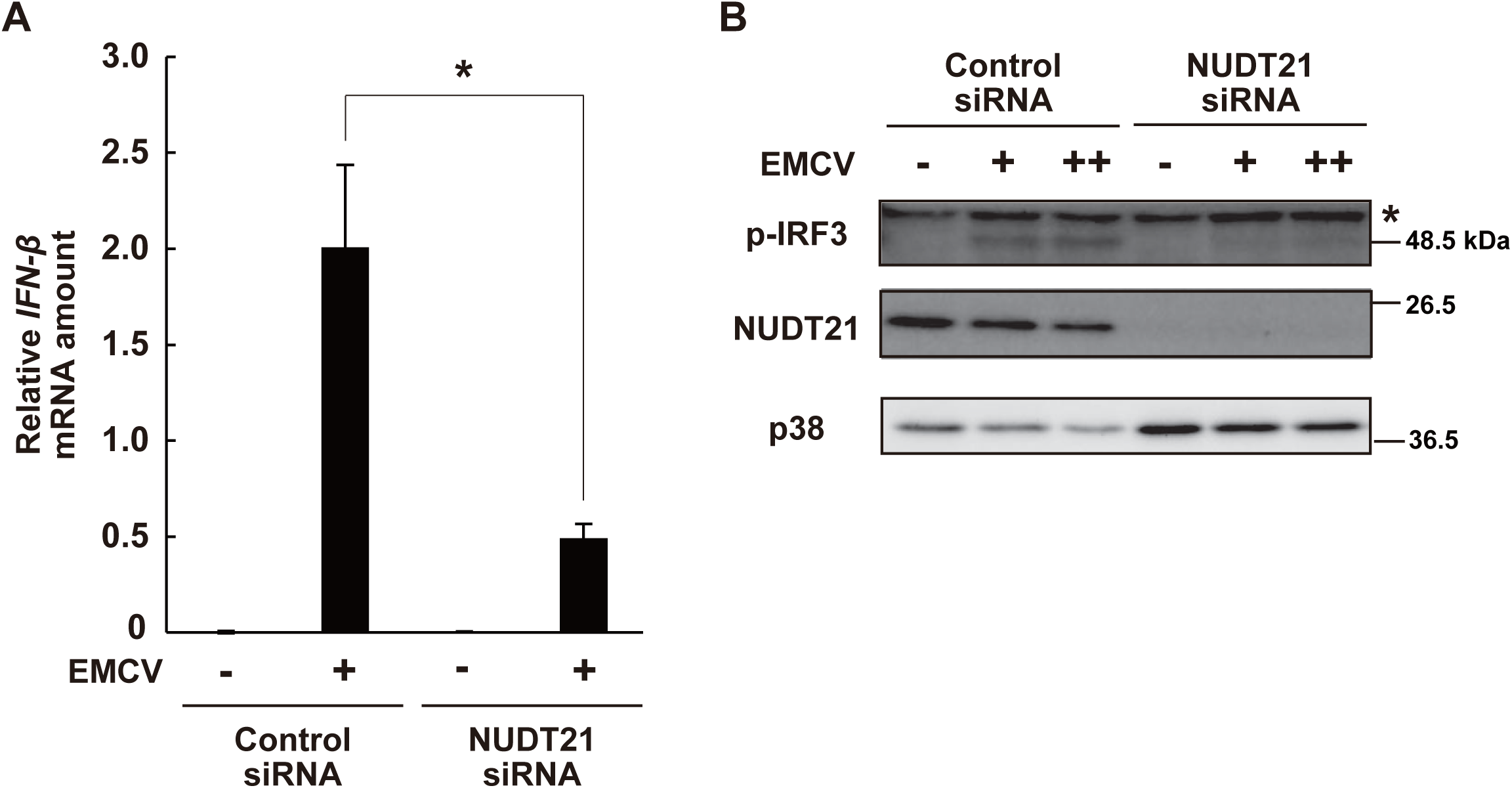
NUDT21 is required for the optimal induction of type I IFN in response to EMCV or NDV infection. (A) Quantitative RT-PCR analysis of IFN-β mRNA in HeLa S3 cells transiently expressing control or NUDT21 siRNAs at 12 h after infection with 0.2 plaque-forming units (PFU) of EMCV. Data are means ± s.e.m. from four independent experiments. \**P* < 0.05 (Student’s *t* test). (B) Immunoblot analysis of phosphorylated IRF3, NUDT21, and p38 in control or NUDT21-knockdown HeLa S3 cells infected with 0.1 (+) or 0.2 (++) PFU of EMCV for 12 h. The asterisk indicates nonspecific bands. Data are representative of three independent experiments.

### Forced localization of IPS-1 to SGs promotes type I IFN induction in response to cytoplasmic dsRNA

Disruption of SGs has been shown to attenuate IFN induction by dsRNA (25, 30), although SGs appear to be dispensable for antiviral responses in some instances (49). We therefore investigated whether forced targeting of IPS-1 to SGs might enhance IFN induction in poly(I:C)-transfected cells. We first constructed an expression vector for a fusion protein containing the SG protein G3BP1 and Myc-tagged IPS-1. However, expression of G3BP1-Myc-IPS-1 resulted in formation of abnormal aggregates within HeLa S3 cells that appeared to include mitochondria as shown by TOMM20 immunostaining (Figure S5). To avoid such mitochondrial aggregation, we generated a fusion protein containing G3BP1 and a Myc-tagged form of IPS-1 that lacks the COOH-terminal TM domain (Figure 5A). We found that this G3BP1-Myc-IPS-1(ΔTM) fusion protein was present mostly in the cytosol of HeLa S3 cells in the absence of poly(I:C) transfection, but preferentially localized to SGs in the presence of poly(I:C) (Figure 5B). We then measured the level of IFN-β mRNA in cells expressing G3BP1, Myc-tagged IPS-1(ΔTM), or G3BP1-Myc-IPS-1(ΔTM) at 9 h after poly(I:C) transfection (Figure 5C, D). Forced expression of G3BP1 resulted in slight enhancement of the increase in the amount of IFN-β mRNA induced by poly(I:C) transfection, consistent with a previous observation (50). We also found that expression of Myc-tagged IPS-1(ΔTM) slightly enhanced the increase in IFN-β mRNA abundance induced by poly(I:C). Expression of G3BP1-Myc-IPS-1(ΔTM) resulted in a markedly greater increase in the level of IFN-β mRNA in poly(I:C)-transfected cells compared with that induced by G3BP1 or Myc-tagged IPS-1(ΔTM) (Figure 5D). Importantly, the amount of G3BP1-Myc-IPS-1(ΔTM) in the cells was markedly lower than that of Myc– IPS(1ΔTM). These results together suggested that the localization of IPS-1 to SGs enhances IFN induction in the presence of cytoplasmic dsRNA.

**Figure 5.**
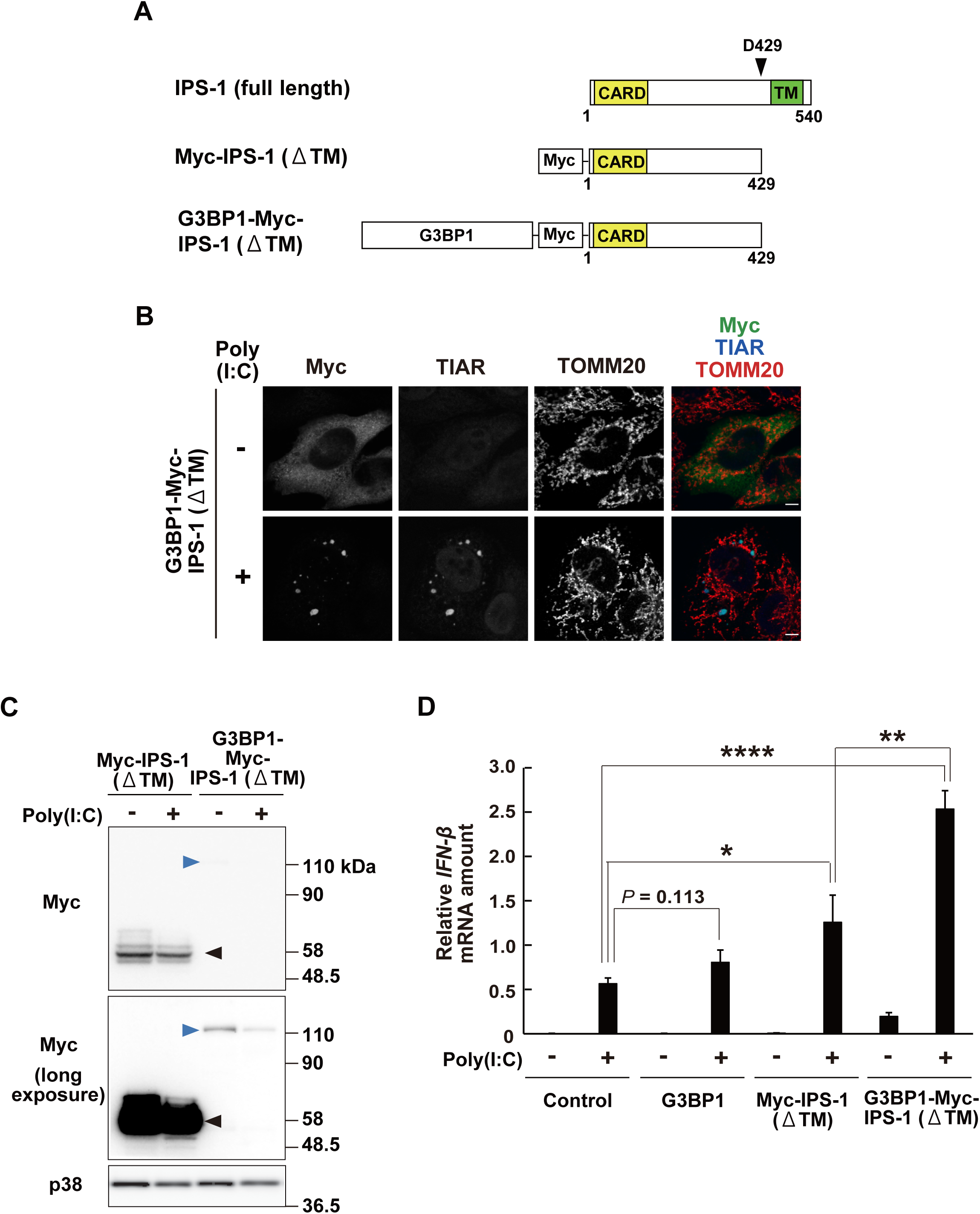
Forced localization of IPS-1 to SGs enhances type I IFN induction in response to poly(I:C) transfection. (A) Schematic representation of human IPS-1, Myc-tagged IPS-1(ΔTM), and G3BP1-Myc-IPS-1(ΔTM). (B) Immunofluorescence analysis of HeLa S3 cells transiently expressing G3BP1-Myc-IPS-1(ΔTM) at 6 h after transfection with poly(I:C) (0.25 μg/ml) or mock transfection. The cells were stained with antibodies to Myc, to TIAR, and to TOMM20. Scale bars, 5 μm. Data are representative of three independent experiments. (C) Immunoblot analysis with antibodies to Myc of HeLa S3 cells transiently expressing Myc–IPS-1(ΔTM) or G3BP1-Myc-IPS-1(ΔTM) at 9 h after transfection with poly(I:C) (0.25 μg/ml) or mock transfection. Black and blue arrowheads indicate Myc– IPS-1(ΔTM) and G3BP1-Myc-IPS-1(ΔTM), respectively. Data are representative of three independent experiments. (D) Quantitative RT-PCR analysis of IFN-β mRNA in HeLa S3 cells transiently expressing G3BP1, Myc–IPS-1(ΔTM), or G3BP1-Myc-IPS-1(ΔTM) at 9 h after transfection with poly(I:C) (0.25 μg/ml) or mock transfection. Data are means ± s.e.m. from four independent experiments. \**P* < 0.05, \*\**P* < 0.01, \*\*\*\**P* < 0.001 (Student’s *t* test).

## Discussion

In the innate immune system, the site of receptor-ligand interaction is often dissociated from that of signal transduction, likely in order to prevent aberrant activation of antipathogen programs in the absence of infection (51). However, these distinct compartments must encounter each other soon after ligand detection. We have now identified NUDT21 as a link between RLR-containing stress granules and mitochondrial IPS-1 as well as an essential mediator of antiviral responses. Our study has therefore unveiled the existence of a cellular machinery that links the site of pathogen recognition and that of antiviral signal transduction.

What is the mechanism by which NUDT21 mediates type I IFN production? Although we cannot exclude the possibility that NUDT21 regulates type I IFN expression through alternative polyadenylation in the nucleus, we propose a model whereby NUDT21 promotes antiviral responses by recruiting IPS-1 to antiviral SGs on the basis of the following observations: (1) Both mass spectrometry and co-immunoprecipitation analyses indicated that NUDT21 forms a complex with mitochondrial IPS-1. (2) A fraction of NUDT21 was found to be localized to mitochondria in resting cells and became localized to SGs in response to poly(I:C) transfection. (3) Another component of the CFIm complex, CFIm68, appeared to be localized only in the nucleus, not being detected in the cytoplasm, suggesting that NUDT21 has a cytoplasmic function independent of the CFIm complex. (4) Knockdown of NUDT21 suppressed the change in the localization of IPS-1 from mitochondria to SGs, but not the formation of SGs, in response to poly(I:C) stimulation. Further studies are required to understand in more detail how NUDT21 regulates IPS-1 localization in response to poly(I:C) transfection.

Caspase-mediated IPS-1 cleavage at a juxtamembrane site has been proposed to inactivate the function of IPS-1 (33, 34). However, an IPS-1 mutant lacking the TM domain was found to form prionlike fibers (“seeds”) that could convert native IPS-1 into functional aggregates (11). The role of IPS-1 cleavage therefore remains controversial. We found that expression of an IPS-1(ΔTM) mutant significantly enhanced the activity of the IFN-β gene promoter only in the presence of full-length IPS-1 (Figure S6). Moreover, our results showed that forced localization of IPS-1(ΔTM) to SGs enhanced type I IFN induction by poly(I:C) transfection. On the basis of these observations, we propose a two-step model: In the early stage of virus infection, the caspase-cleaved form of IPS-1 cooperates with full-length IPS-1 to form large aggregates on the mitochondrial surface that associate with RLRs in SGs and thereby elicit antiviral responses. In the late stage, caspases cleave the remaining intact IPS-1 molecules and thereby terminate IPS-1–mediated immune responses. This model may reconcile the apparent discrepancy in the effects of caspase-mediated cleavage on IPS-1 function mentioned above.

The precise nature of the physical interaction between SGs and IPS-1 aggregates remains unclear. Antiviral SGs are membraneless organelles that are formed by liquid-liquid phase separation within the cytoplasm. We observed that a fraction of IPS-1 appeared to form fiberlike structures rather than being evenly distributed within SGs (Figure 2A). This observation and the previous findings on the prionlike aggregation of IPS-1 (11) suggest that IPS-1 aggregates may form a subcompartment within SGs, perhaps by undergoing a liquid-to-solid phase transition (which typically underlies the formation of solid prionlike aggregates or crystals), or that they may lie adjacent to SGs and associate with RIG-I in SGs at the interface. In these two cases, the presence of NUDT21 may facilitate functional solidification of IPS-1 in SGs or functional association between RIG-I in SGs and IPS-1 aggregates, respectively.

Lysine-63–linked polyubiquitination by the ubiquitin ligase TRIM25 has been implicated in the activation of RIG-I and subsequent formation of large IPS-1 aggregates (52, 53, 11). It would thus be of interest to examine the relation between such TRIM25-mediated polyubiquitination and NUDT21 in the formation of IPS-1 aggregates and their association with SGs. Of note, RIG-I has been shown to associate exclusively with either TRIM25 or IPS-1, with the two complexes being localized to distinct compartments (54). Together with our observation that IPS-1 is not distributed among all SGs in a cell, this finding suggests that RIG-I might shift from the TRIM25 compartment to the IPS-1 compartment in association with IPS-1 activation (54) and that NUDT21 may facilitate this transition.

In closing, we have here demonstrated the cytoplasmic localization of NUDT21 and its unexpected role in regulation of antiviral proteins in the cytoplasm in addition to its well-established localization to nucleus and function in alternative polyadenylation. We therefore propose that NUDT21 may function in broader biological contexts, at least at mitochondria and SGs, than anticipated, and our results provide a basis for the development of a new target for clinical intervention in viral propagation.

## Acknowledgments

We thank M. Okajima and K. Takechi (Graduate School of Pharmaceutical Sciences, The University of Tokyo) for technical assistance; M. Yoneyama (Chiba University) for providing the pEF-BOS-FLAG-IPS-1 plasmid, antibodies to RIG-I, and ECMV; and laboratory members for discussion.

## Author Contributions

S.A.I. performed experiments and analyzed data. T.N. and S.I. conducted the mass spectrometric analysis. Y.O. provided the pN1/G3BP1-iRFP plasmid and technical assistance for imaging analysis. S.A.I., Y.G., and T.O. conceived the study and wrote the manuscript.

## References

1. Stetson, D.B., and R. Medzhitov. 2006. Type I interferons in host defense. Immunity. 25: 373–381.

2. Schneider, W.M., M.D. Chevillotte, and C.M. Rice. 2014. Interferon-stimulated genes: a complex web of host defenses. Annu. Rev. Immunol. 32: 513–545.

3. Akira, S., S. Uematsu, and O. Takeuchi. 2006. Pathogen recognition and innate immunity. Cell. 124: 783–801.

4. Yoneyama, M., M. Kikuchi, T. Natsukawa, N. Shinobu, T. Imaizumi, M. Miyagishi, K. Taira, S. Akira, and T. Fujita. 2004. The RNA helicase RIG-I has an essential function in double-stranded RNA-induced innate antiviral responses. Nat Immunol. 5: 730–737.

5. Gitlin, L., W. Barchet, S. Gilfillan, M. Cella, B. Beutler, R.A. Flavell, M.S. Diamond, and M. Colonna. 2006. Essential role of mda-5 in type I IFN responses to polyriboinosinic: polyribocytidylic acid and encephalomyocarditis picornavirus. Proc Natl Acad Sci USA. 103: 8459–8464.

6. Kato, H., O. Takeuchi, S. Sato, M. Yoneyama, M. Yamamoto, K. Matsui, S. Uematsu, A. Jung, T. Kawai, K.J. Ishii, O. Yamaguchi, K. Otsu, T. Tsujimura, C.S. Koh, C. Reis e Sousa, Y. Matsuura, T. Fujita, and S. Akira. 2006. Differential roles of MDA5 and RIG-I helicases in the recognition of RNA viruses. Nature. 441: 101–105.

7. Kawai, T., K. Takahashi, S. Sato, C. Coban, H. Kumar, H. Kato, K.J. Ishii, O. Takeuchi, and S. Akira. 2005. IPS-1, an adaptor triggering RIG-I- and Mda5-mediated type I interferon induction. Nat Immunol. 6: 981–988.

8. Seth, R.B., L. Sun, C.K. Ea, and Z.J. Chen. 2005. Identification and characterization of MAVS, a mitochondrial antiviral signaling protein that activates NF-κB and IRF3. Cell. 122: 669–682.

9. Meylan, E., J. Curran, K. Hofmann, D. Moradpour, M. Binder, R. Bartenschlager, and J. Tschopp. 2005. Cardif is an adaptor protein in the RIG-I antiviral pathway and is targeted by hepatitis C virus. Nature. 437: 1167–1172.

10. Xu, L.G., Y.Y. Wang, K.J. Han, L.Y. Li, Z. Zhai, and H.B. Shu. 2005. VISA is an adapter protein required for virus-triggered IFN-β signaling. Mol Cell. 19: 727–740.

11. Hou, F., L. Sun, H. Zheng, B. Skaug, Q.X. Jiang, and Z.J. Chen. 2011. MAVS forms functional prion-like aggregates to activate and propagate antiviral innate immune response. Cell. 146: 448–461.

12. Fitzgerald, K.A., S.M. Mcwhirter, K.L. Faia, D.C. Rowe, E. Latz, D.T. Golenbock, A.J. Coyle, S.M. Liao, and T. Maniatis. 2003. IKKε and TBK1 are essential components of the IRF3 signaling pathway. Nat Immunol. 4: 491–496.

13. Chu, W.M., D. Ostertag, Z.W. Li, L. Chang, Y. Chen, Y. Hu, B. Williams, J. Perrault, and M. Karin. 1999. JNK2 and IKKβ are required for activating the innate response to viral infection. Immunity. 11: 721–731.

14. Du, W., and T. Maniatis. 1992. An ATF / CREB binding site is required for virus induction of the human interferon beta gene. Proc Natl Acad Sci USA. 89: 2150–2154.

15. Honda, K., A. Takaoka, and T. Taniguchi. 2006. Type I inteferon gene induction by the interferon regulatory factor family of transcription factors. Immunity. 25: 349–360.

16. Okazaki, T., M. Higuchi, K. Takeda, K. Iwatsuki-Horimoto, M. Kiso, M. Miyagishi, H. Yanai, A. Kato, M. Yoneyama, T. Fujita, T. Taniguchi, Y. Kawaoka, H. Ichijo, and Y. Gotoh. 2015. The ASK family kinases differentially mediate induction of type I interferon and apoptosis during the antiviral response. Sci Signal. 8: ra78.

17. Kumar, H., T. Kawai, H. Kato, S. Sato, K. Takahashi, C. Coban, M. Yamamoto, S. Uematsu, K.J. Ishii, O. Takeuchi, and S. Akira. 2006. Essential role of IPS-1 in innate immune responses against RNA viruses. J Exp Med. 203: 1795–1803.

18. Sun, Q., L. Sun, H.H. Liu, X. Chen, R.B. Seth, J. Forman, and Z.J. Chen. 2006. The specific and essential role of MAVS in antiviral innate immune responses. Immunity. 24: 633–642.

19. Yoneyama, M., M. Kikuchi, K. Matsumoto, T. Imaizumi, M. Miyagishi, K. Taira, E. Foy, Y.M. Loo, M. Gale Jr., S. Akira, S. Yonehara, A. Kato, and T. Fujita. 2005. Shared and unique functions of the DExD/H-Box helicases RIG-I, MDA5, and LGP2 in antiviral innate immunity. J Immunol. 175: 2851–2858.

20. Buchan, J.R., and R. Parker. 2009. Eukaryotic stress granules : the ins and outs of translation. Mol Cell. 36: 932–941.

21. McCormick, C., and D.A. Khaperskyy. 2017. Translation inhibition and stress granules in the antiviral immune response. Nat Rev Immunol. 17: 647–660.

22. Kedersha, N.L., M. Gupta, W. Li, I. Miller, and P. Anderson. 1999. RNA-binding proteins TIA-1 and TIAR link the phosphorylation of eIF-2α to the assembly of mammalian stress granules. J Cell Biol. 147: 1431–1442.

23. Tourrière, H., K. Chebli, L. Zekri, B. Courselaud, J.M. Blanchard, E. and Bertrand, J. Tazi. 2003. The RasGAP-associated endoribonuclease G3BP assembles stress granules. J Cell Biol. 160: 823–831.

24. Boeynaems, S., S. Alberti, N.L. Fawzi, T. Mittag, M. Polymenidou, F. Rousseau, J. Schymkowitz, J. Shorter, B. Wolozin, L. Van Den Bosch, P. Tompa, and M. Fuxreiter. 2018. Protein Phase Sparation: A New Phase in Cell Biology. Trends Cell Biol. 28: 420–435.

25. Onomoto, K., M. Jogi, J.S. Yoo, R. Narita, S. Morimoto, A. Takemura, S. Sambhara, A. Kawaguchi, S. Osari, K. Nagata, T. Matsumiya, H. Namiki, M. Yoneyama, and T. Fujita. 2012. Critical role of an antiviral stress granule containing RIG-I and PKR in viral detection and innate immunity. PLoS One. 7: e43031.

26. Oh, S.W., K. Onomoto, M. Wakimoto, K. Onoguchi, F. Ishidate, T. Fujiwara, M. Yoneyama, H. Kato, and T. Fujita. 2016. Leader-Containing Uncapped Viral Transcript Activates RIG-I in Antiviral Stress Granules. PLoS Pathog. 12: e1005444.

27. White, J.P., A.M. Cardenas, W.E. Marissen, and R.E. Lloyd. 2007. Inhibition of cytoplasmic mRNA stress granule formation by a viral proteinase. Cell Host Microbe. 2: 295–305.

28. Khaperskyy, D.A., T.F. Hatchette, and C. McCormick. 2012. Influenza A virus inhibits cytoplasmic stress granule formation. FASEB J. 26: 1629–1639.

29. Borghese, F., and T. Michiels. 2011. The leader protein of cardioviruses inhibits stress granule assembly. J Virol. 85: 9614–9622.

30. Ng, C.S., M. Jogi, J.S. Yoo, K. Onomoto, S. Koike, T. Iwasaki, M. Yoneyama, H. Kato, and T. Fujita. 2013. Encephalomyocarditis virus disrupts stress granules, the critical platform for triggering antiviral innate immune responses. J Virol. 87: 9511–9522.

31. Finnen, R.L., M. Zhu, J. Li, D. Romo, and B.W. Banfield. 2016. Herpes Simplex Virus 2 Virion Host Shutoff Endoribonuclease Activity Is Required To Disrupt Stress Granule Formation. J Virol. 90: 7943–7955.

32. Li, X.D., L. Sun, R.B. Seth, G. Pineda, and Z.J. Chen. 2005. Hepatitis C virus protease NS3/4A cleaves mitochondrial antiviral signaling protein off the mitochondria to evade innate immunity. Proc Natl Acad Sci USA. 102: 17717–17722.

33. Rebsamen, M., E. Meylan, J. Curran, and J. Tschopp. 2008. The antiviral adaptor proteins Cardif and Trif are processed and inactivated by caspases. Cell Death Differ. 15: 1804–1811.

34. Ning, X., Y. Wang, M. Jing, M. Sha, M. Lv, P. Gao, R. Zhang, X. Huang, J.M. Feng, and Z. Jiang. 2019. Apoptotic Caspases Suppress Type I Interferon Production via the Cleavage of cGAS, MAVS, and IRF3. Mol Cell. 74: 19-31.e7.

35. Zhang, Peifen, Y. Li, J. Xia, J. He, J. Pu, J. Xie, S. Wu, L. Feng, X. Huang, and Ping Zhang. 2014. IPS-1 plays an essential role in dsRNA-induced stress granule formation by interacting with PKR and promoting its activation. J Cell Sci. 127: 2471–2482.

36. Brown, K.M., and G.M. Gilmartin. 2003. A mechanism for the regulation of pre-mRNA 3’ processing by human cleavage factor I m. Mol Cell. 12: 1467–1476.

37. Di Giammartino, D.C., K. Nishida, and J.L. Manley. 2011. Mechanisms and consequences of alternative polyadenylation. Mol Cell. 43: 853–866.

38. Yang, Q., M. Coseno, G.M. Gilmartin, and S. Doublié. 2011. Crystal structure of a human cleavage factor CFI(m)25 / CFI(m)68 / RNA complex provides an insight into poly (A) site recognition and RNA looping. Structure. 19: 368–377.

39. Elkon, R., A.P. Ugalde, and R. Agami. 2013. Alternative cleavage and polyadenylation: extent, regulation and function. Nat Rev Genet. 14: 496–506.

40. Masamha, C.P., Z. Xia, J. Yang, T.R. Albrecht, M. Li, A.B. Shyu, W. Li, and E.J. Wagner. 2014. CFIm25 links alternative polyadenylation to glioblastoma tumour suppression. Nature. 510: 412–416.

41. Brumbaugh, J., B. Di Stefano, X. Wang, M. Borkent, E. Forouzmand, K.J. Clowers, F. Ji, B.A. Schwarz, M. Kalocsay, S.J. Elledge, Y. Chen, R.I. Sadreyev, S.P. Gygi, G. Hu, Y. Shi, and K. Hochedlinger. 2018. Nudt21 Controls Cell Fate by Connecting Alternative Polyadenylation to Chromatin Signaling. Cell. 172: 106-120.e21.

42. Jia, X., S. Yuan, Y. Wang, Y. Fu, Yong Ge, Yutong Ge, X. Lan, Y. Feng, F. Qiu, P. Li, S. Chen, and A. Xu. 2017. The role of alternative polyadenylation in the antiviral innate immune response. Nat Commun. 8: 14605.

43. Gaucherand, L., B.K. Porter, R.E. Levene, E.L. Price, S.K. Schmaling, C.H. Rycroft, Y. Kevorkian, C. McCormick, D.A. Khaperskyy, and M.M. Gaglia. 2019. The Influenza A Virus Endoribonuclease PA-X Usurps Host mRNA Processing Machinery to Limit Host Gene Expression. Cell Rep. 27: 776-792.e7.

44. Okazaki, T., M. Higuchi, and Y. Gotoh. 2013. Mitochondrial localization of the antiviral signaling adaptor IPS-1 is important for its induction of caspase activation. Genes Cells. 18: 493–501.

45. Natsume, T., Y. Yamauchi, H. Nakayama, T. Shinkawa, M. Yanagida, N. Takahashi, and T. Isobe. 2002. A direct nanoflow liquid chromatography - tandem mass spectrometry system for interaction proteomics. Anal Chem. 74: 4725–4733.

46. Costes, S.V., D. Daelemans, E.H. Cho, Z. Dobbin, G. Pavlakis, and S. Lockett. 2004. Automatic and quantitative measurement of protein-protein colocalization in live cells. Biophys J. 86: 3993–4003.

47. Manders, E.M.M., F.J. Verbeek, and J.A. Aten. 1993. Measurement of colocalization of objects in dual-colour confocal images. J. Microsc. 169: 375–382.

48. Mikkelsen, S.S., S.B. Jensen, S. Chiliveru, J. Melchjorsen, I. Julkunen, M. Gaestel, J.S. Arthur, R.A. Flavell, S. Ghosh, and S.R. Paludan. 2009. RIG-I-mediated activation of p38 MAPK is essential for viral induction of interferon and activation of dendritic cells: dependence on TRAF2 and TAK1. J Biol Chem. 284: 10774–10782.

49. Langereis, M.A., Q. Feng, and F.J. van Kuppeveld. 2013. MDA5 localizes to stress granules, but this localization is not required for the induction of type I interferon. J Virol. 87: 6314–6325.

50. Kim, S.S., L. Sze, C. Liu, and K.P. Lam. 2019. The stress granule protein G3BP1 binds viral dsRNA and RIG-I to enhance interferon-β response. J Biol Chem. 294: 6430–6438.

51. Kagan, J.C. 2012. Signaling organelles of the innate immune system. Cell. 151: 1168–1178..

52. Gack, M.U., Y.C. Shin, C.H. Joo, T. Urano, C. Liang, L. Sun, O. Takeuchi, S. Akira, Z. Chen, S. Inoue, and J.U. Jung. 2007. TRIM25 RING-finger E3 ubiquitin ligase is essential for RIG-I-mediated antiviral activity. Nature. 446: 916–920.

53. Jiang, X., L.N. Kinch, C.A. Brautigam, X. Chen, F. Du, N.V. Grishin, and Z.J. Chen. 2012. Ubiquitin-induced oligomerization of the RNA sensors RIG-I and MDA5 activates antiviral innate immune response. Immunity. 36: 959–973.

54. Sánchez-Aparicio, M.T., J. Ayllón, A. Leo-Macias, T. Wolff, and A. García-Sastre. 2017. Subcellular Localizations of RIG-I, TRIM25, and MAVS Complexes. J Virol. 91: e01155–16.

