## Supplemental Figure1-6 for "NUDT21 links mitochondrial IPS-1 to RLR-containing stress granules and activates host antiviral defense"

### Supplemental Figure 1

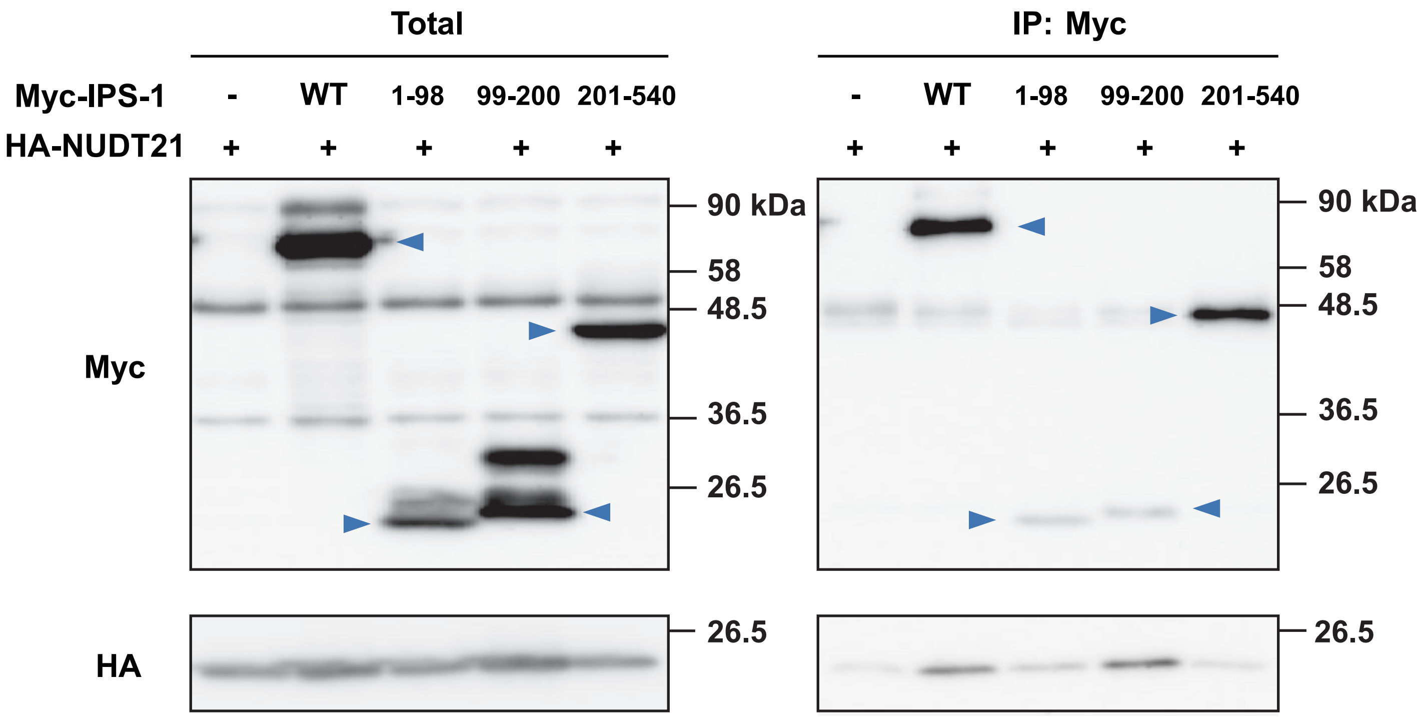


**Figure S1. NUDT21 forms a complex with IPS-1 fragments including CARD or the proline-rich domain.**

Extracts of HEK293T cells transiently expressing HA-tagged NUDT21 and Myc-tagged full-length (WT) IPS-1 or fragments thereof (amino acids 1–98, 99–200, or 201–540) were subjected to immunoprecipitation (IP) with antibodies to Myc, and the resulting precipitates as well as the original cell extracts (Total) were subjected to immunoblot analysis with antibodies to Myc and to HA. Arrowheads indicate each IPS-1 protein. Data are representative of three independent experiments.

#### Supplemental Figure 2

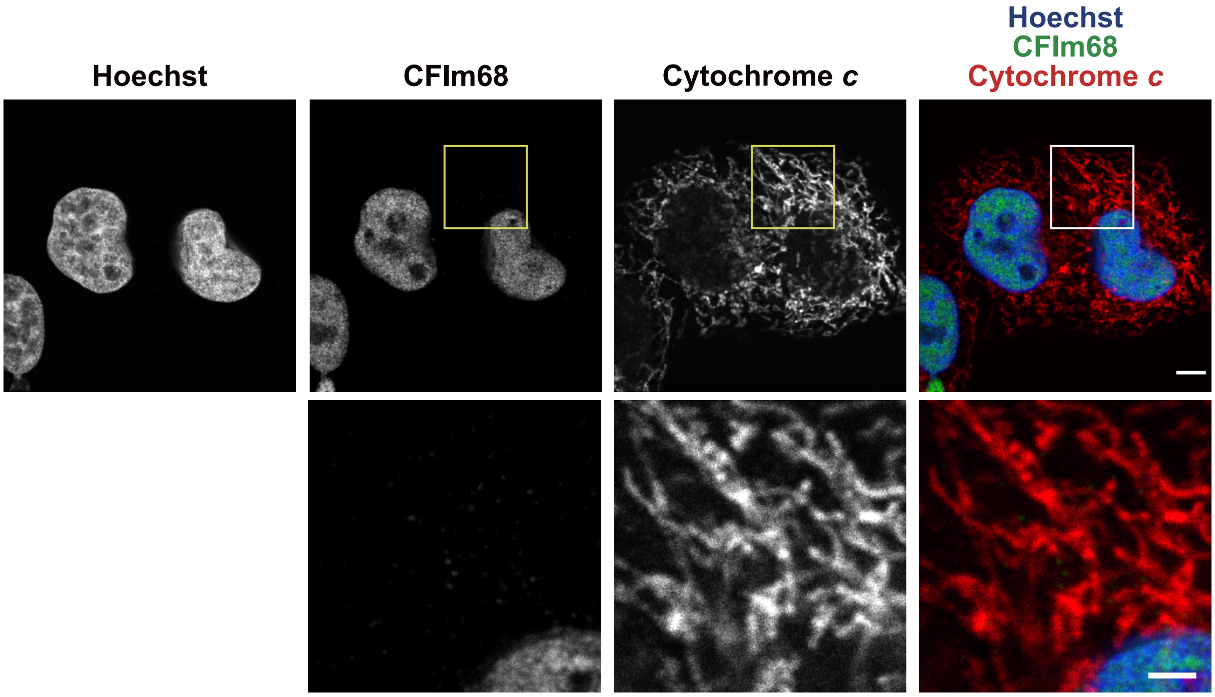


**Figure S2. CFIm68 localizes to the nucleus but not to mitochondria.**
HeLa S3 cells were subjected to immunofluorescence staining with antibodies to CFIm68 and to cytochrome c. Nuclei were stained with Hoechst 33342. The boxed regions of the upper panels are shown at higher magnification in the corresponding lower panels. Scale bars, 5 μm or 2 μm (higher magnification images). Data are representative of three independent experiments.

#### Supplemental Figure 3

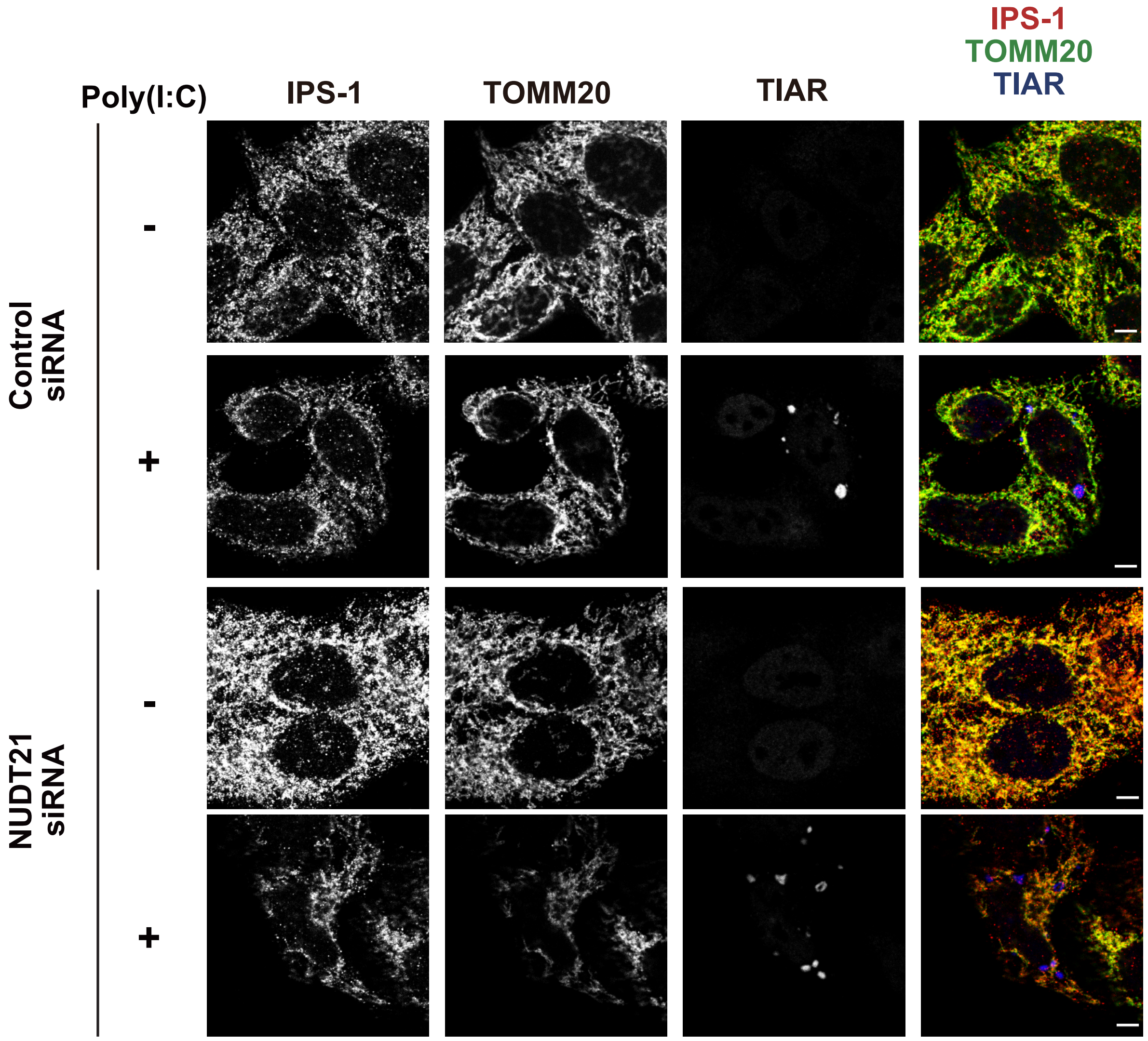


**Figure S3. NUDT21 knockdown does not affect the distribution of IPS-1 in mock-treated cells but attenuates the colocalization of IPS-1 and TIAR in poly(I:C)-transfected cells.**

HeLa S3 cells expressing control or NUDT21 siRNAs were subjected to immunofluorescence staining with antibodies to IPS-1, to TOMM20, and to TIAR at 6 h after transfection with poly(I:C) (0.25 μg/ml) or mock transfection. Scale bars, 5 μm. Data are representative of three independent experiments.

### Supplemental Figure 4

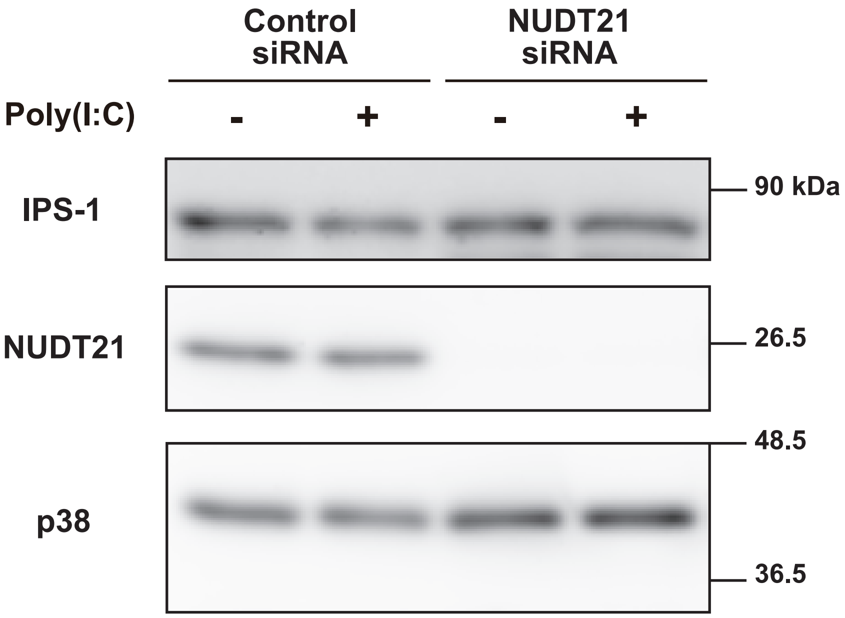


**Figure S4. NUDT21 knockdown does not affect the expression level of IPS-1.**
HeLa S3 cells expressing control or NUDT21 siRNAs were subjected to immunoblot analysis with antibodies to IPS-1, to NUDT21, and to p38 (loading control) at 9 h after transfection with poly(I:C) (0.25 μg/ml) or mock transfection. Data are representative of three independent experiments.

#### Supplemental Figure 5

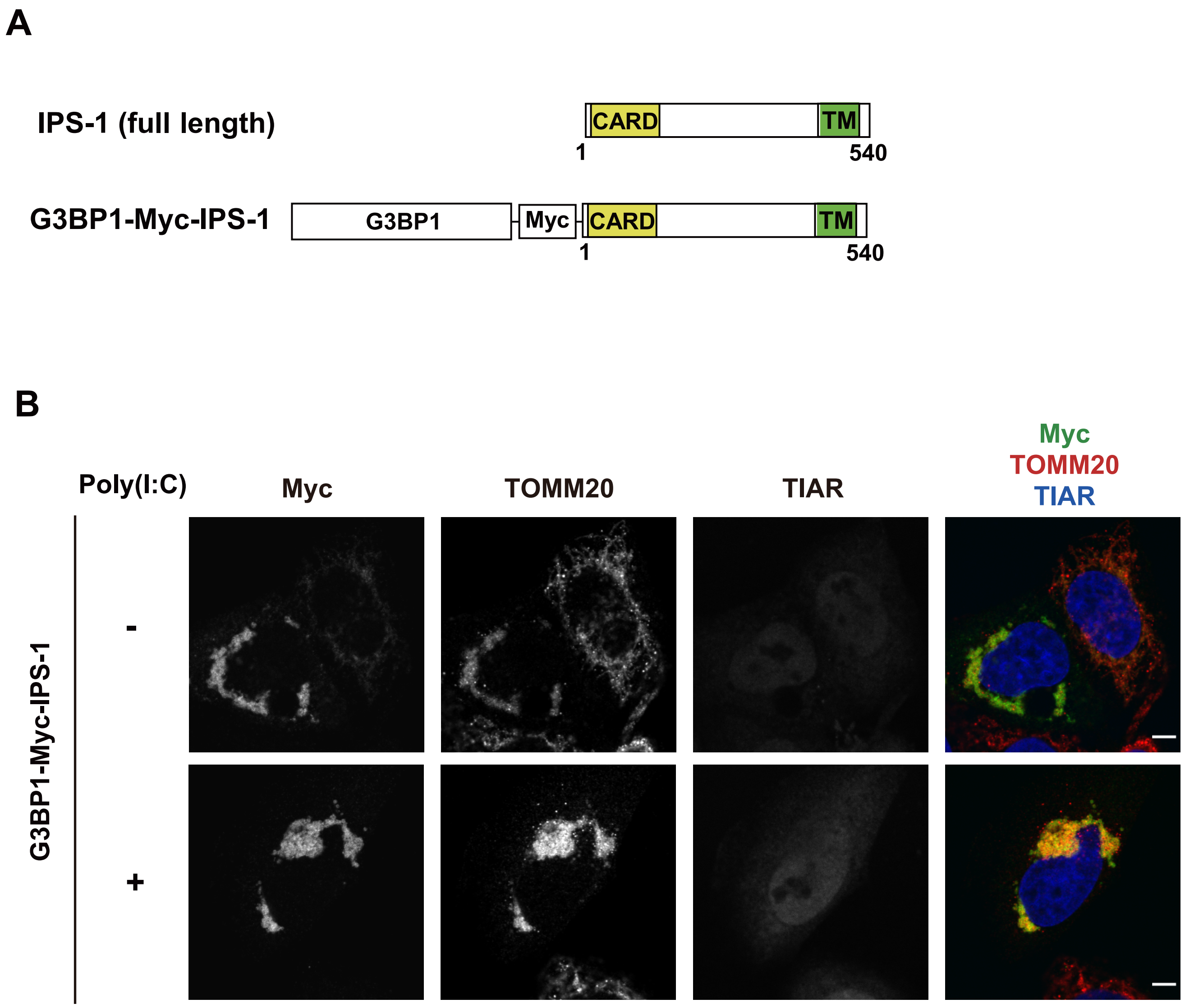


**Figure S5. Predominant localization of the G3BP1-Myc-IPS-1 fusion protein to mitochondria.**

(A) Schematic representation of human IPS-1 and G3BP1-Myc-IPS-1.

(B) Immunofluorescence analysis of HeLa S3 cells transiently expressing G3BP1-Myc-IPS-1 at 6 h after transfection with poly(I:C) (0.25 μg/ml) or mock transfection. The cells were stained with antibodies to Myc, to TOMM20, and to TIAR. Scale bars, 5 μm. Data are representative of three independent experiments.

#### Supplemental Figure 6

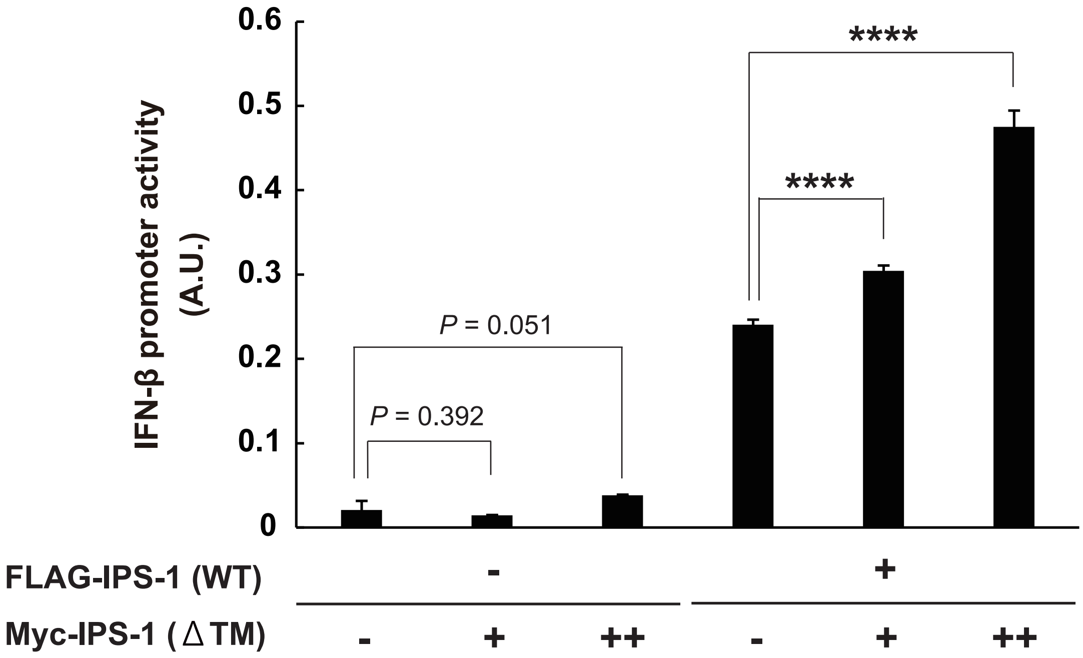


**Figure S6. Expression of IPS-1(ΔTM) promotes type I IFN induction by full-length IPS-1.**

HEK293T cells were transiently transfected with a luciferase reporter plasmid containing the IFN-β gene promoter as well as with 5 ng of an expression plasmid encoding FLAG-tagged full-length IPS-1 and 50 ng (+) or 500 ng (++) of a plasmid for Myc-tagged IPS-1(ΔTM). The cells were subsequently assayed for luciferase activity as a measure of IFN-β gene promoter activity. Data are presented in arbitrary units (A.U.) and are means ± s.d. of triplicates from an experiment that was repeated a total of three times with similar results. ****$P < 0.001$ (Student's *t* test).
